# Distinct regulation of AEBP2 isoforms on PRC2 activity

**DOI:** 10.1101/2025.11.09.687442

**Authors:** Yingying Li, Hanbyeol Kim, Chul-Hwan Lee

**Affiliations:** Department of Pharmacology, Seoul National University College of Medicine, Seoul, Republic of Korea; Department of Biomedical Sciences, Seoul National University College of Medicine, Seoul, Republic of Korea; Cancer Research Institute, Ischemic/hypoxic Disease Institute, Neuroscience Research Institute, Institute of Molecular Biology & Genetics, Seoul National University, Seoul, Republic of Korea; Wide River Institute of Immunology, Seoul National University, Hongcheon, Republic of Korea

**Keywords:** AEBP2 long and short isoforms, embryonic stem cell, epigenetic regulation, histone modification, H3K27me3, PRC2

## Abstract

Polycomb Repressive Complex 2 (PRC2) represses gene expression through catalyzing H3K27me3, a histone modification essential for gene repression and maintenance of cellular identity. PRC2 accessory proteins modulate its catalytic activity, chromatin localization, and propagation along the chromatin. AEBP2, one of the PRC2 accessory proteins required for mouse embryogenesis, exists in distinct isoforms. The short isoform enhances PRC2 catalytic activity and promotes H3K27me3 spreading, facilitating robust gene repression while the function of the long isoform has remained unclear. Here, we found that the N-terminal region of AEBP2 long isoform contains a conserved DE-rich motif that inhibits PRC2 activity, both EZH2 automethylation and H3K27 methylation. In addition, re-expression of the AEBP2 long isoform in *Mtf2*/*Jarid2*/*Aebp2* triple-knockout mESCs failed to restore H3K27me3 and caused defective differentiation, unlike the short isoform. Together, our findings uncover an isoform-specific regulatory mechanism by which AEBP2 controls PRC2 activity and contribute to a broader understanding of how PRC2 is dynamically regulated during development.

## Introduction

Polycomb Repressive Complex 2 (PRC2) is a key epigenetic regulator that represses the expression of lineage-specific genes (Bracken *et al*, 2006; Barski *et al*, 2007). PRC2 achieves gene silencing by catalyzing mono-, di-, and tri-methylation on histone H3 lysine 27, and H3K27me3 is a hallmark of facultative heterochromatin (Lee *et al*, 2022; Yu *et al*, 2019). PRC2 comprises three core subunits: EZH1 or EZH2, EED, and SUZ12. EZH1 and EZH2 are the catalytic subunits of PRC2 that harbor SET domains, EED contains an aromatic cage that can bind to H3K27me3 and allosterically activate PRC2, SUZ12 functions as a scaffold protein which interacts with other core subunits and its accessory proteins (Lee *et al*, 2018c; Margueron *et al*, 2009; Yu *et al*, 2019; Lee *et al*, 2018a; Li *et al*, 2017; Oksuz *et al*, 2018; Chen *et al*, 2018; Son *et al*, 2013; Kasinath *et al*, 2018; Lee *et al*, 2018b). Additionally, RBAP48 associates with the PRC2 core subunits to enhance its functionality. Besides the core subunits, PRC2 accessory proteins, including AEBP2, JARID2, Polycomb-like proteins (MTF2, PHF1 and PHF19), EPOP and PALI associate with SUZ12, are critical factors to recruit PRC2 to chromatin and enhance its catalytic activity on H3K27 (Brien *et al*, 2012; Gatchalian *et al*, 2015; Zhang *et al*, 2011b; Højfeldt *et al*, 2019a; Kasinath *et al*, 2021; Lee *et al*, 2018a; Zhang *et al*, 2021; Kasinath *et al*, 2018). Based on their accessory components, PRC2 complexes are classified into two major subtypes: PRC2.1, which includes either PHF1, MTF2, or PHF19 along with PALI or EPOP; and PRC2.2, which contains JARID2 and AEBP2. Notably, MTF2 and JARID2 are two major proteins indispensable for the initial recruitment of PRC2 at CpG island containing GCN- or GA-rich sequence in mouse embryonic stem cells (mESCs) (Healy *et al*, 2019; Højfeldt *et al*, 2019b; Perino *et al*, 2018; Conway *et al*, 2018; Sarma *et al*, 2008; Li *et al*, 2010; Wang *et al*, 2017; Kasinath *et al*, 2021; Cooper *et al*, 2016; Choi *et al*, 2017).

The presence and expression of different PRC2 accessory proteins are critical for precise control of PRC2 activity in distinct cellular contexts and at specific developmental stages. Previous studies revealed that JARID2 containing PRC2.2 subcomplex is involved in specific gene regulatory processes during development, contributing to *de novo* formation of H3K27me3 repressive chromatin domains, while MTF2 containing PRC2.1 subcomplex is responsible for maintaining pre-existing H3K27me3 repressive domains that are already established in the early embryonic stage (Petracovici & Bonasio, 2021). How other PRC2 accessory proteins spatiotemporally regulate PRC2 recruitment and activity remain unclear.

AEBP2, an accessory protein, plays a vital role in the regulation of PRC2 activity. It facilitates the interaction between JARID2 and SUZ12, promoting the assembly of the PRC2.2 subcomplex (Kasinath *et al*, 2021). Furthermore, a recent structural investigation has shown that both AEBP2 and JARID2 interact independently but synergistically with ubiquitinated H2AK119, a critical histone modification that aids in heterochromatin formation (Kasinath *et al*, 2021). Studies with mice lacking AEBP2 have demonstrated that its deletion results in later-stage embryonic lethality. In addition, mice with one functional copy of AEBP2 (heterozygous deletion) exhibit a variety of abnormalities, including enlarged hepatic veins, jugular lymph node sac, enlarged colon, and hypopigmentation, highlighting the essential role of AEBP2 in organogenesis and tissue specification (Kim *et al*, 2011; Grijzenhout *et al*, 2016).

AEBP2 is predominantly present in two distinct isoforms, long and short isoforms (Kim *et al*, 2011). Previous studies have demonstrated that the short isoform of AEBP2 enhances the activity of PRC2, specifically its histone methyltransferase (HMT) activity, by promoting nucleosome binding (Lee *et al*, 2018a). However, the impact of the long isoform, specifically its basic-rich N-terminal region, on PRC2 activity remains unclear. Interestingly, the AEBP2 long isoform was identified as the predominant form in adult mouse organs, whereas the AEBP2 short isoform was specifically expressed during the embryonic stage (Kim *et al*, 2015). This indicates that the two isoforms may have distinct functions in different cellular contexts or developmental stages.

In this study, we investigate how the two isoforms of AEBP2, defined by the presence or absence of an N-terminal acidic-rich region, differentially regulate PRC2 catalytic activity. We observed that the long isoform of AEBP2, which contains this acidic-rich domain, contributes to the inhibition of PRC2 activity, in sharp contrast to the stimulatory effect exerted by the short isoform. By combining *in vitro* reconstitution with cellular assays, we aim to uncover how these opposing influences fine-tune PRC2 function in a context-dependent manner. This work provides a framework to understand how differential isoform expression of AEBP2 may act as a developmental switch in modulating PRC2-mediated gene repression.

## Results

### The AEBP2 long isoform inhibits while the short isoform stimulates PRC2 activity

AEBP2 long and short isoforms originate from the use of alternative start exons (Fig. S1A-C). To investigate how each isoform differentially modulates PRC2 activity, either the short isoform (human transcript variant 3) or the long isoform (human transcript variant 4) was ectopically expressed in *Aebp2* knockout (KO) mESCs (Fig. 1A, top). The global level of H3K27me3 was marginally reduced when expressing the long isoform (Fig.1B). Since other PRC2 accessory factors might compensate for Aebp2 loss, we extended this analysis using triple-knockout (TKO) mESCs lacking *Mtf2*, *Jarid2*, and *Aebp2* (Fig. 1A, bottom). In this context, reintroducing the AEBP2 long isoform resulted in a marked decrease in H3K27me3 levels (Fig. 1C). These findings suggest that, unlike the previously characterized short isoform (Lee *et al*, 2018a), the long isoform of AEBP2 can act as a negative regulator of PRC2 activity. To directly compare the PRC2 activity between the short and long isoforms of AEBP2, we purified PRC2 reconstituted with either the AEBP2 long or short isoform together with the core subunits and RBAP46/48 (Fig. 1D). The HMT assay using unmodified di-nucleosome showed that the short isoform stimulated PRC2 activity while the long isoform inhibited it (Fig. 1E). This inhibitory effect of the long isoform was also observed when using H2AK119ub-modified mono-nucleosome (Fig. 1F), which is known to interact with AEBP2 through its Zn-finger domain (Kasinath *et al*, 2021).

**Figure 1.**
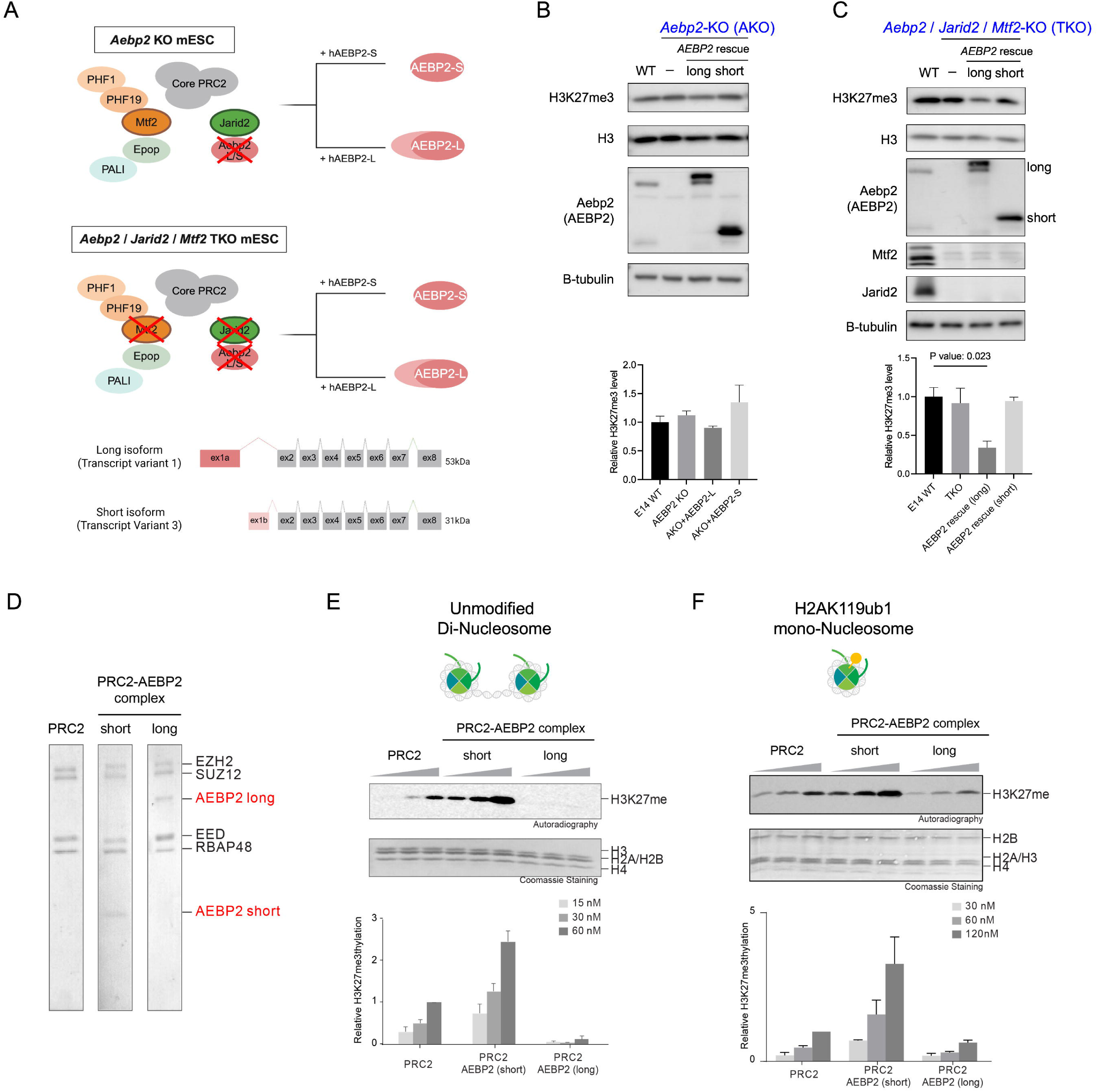
AEBP2 short isoform stimulates PRC2 while AEBP2 long isoform inhibits PRC2 activity. **(A)** Schematic diagram showing the generation of *Aebp2* KO mESCs or *Mtf2*/*Jarid2*/*Aebp2* Triple KO (TKO) cells, followed by rescue with either human AEBP2 short or long isoform (top). Schematic representation of the *AEBP2* gene structure showing alternative transcription start sites that produce two isoforms. The long isoform initiates from exon 1a, while the short isoform originates from exon 1b, resulting in proteins of approximately 53 kDa (AEBP2 long isoform) and 31 kDa (AEBP2 short isoform), respectively (bottom). **(B)** Western blot analysis of H3K27me3, H3, Aebp2 (AEBP2) and b-tubulin in WT mESCs or cells carrying *Aebp2* KO or *Aebp2* KO rescued with specific AEBP2 isoforms (top). Quantification of H3K27me3 levels is shown (n = 3 per data point; mean ± SD) (bottom). **(C)** Western blot analysis of H3K27me3, H3, Aebp2 (AEBP2) and b-tubulin in WT mESCs, *Mtf2*/*Jarid2*/*Aebp2* TKO cells, and TKO cells rescued with specific AEBP2 isoforms (top). Quantification of H3K27me3 levels is shown (n = 3 per data point; mean ± SD) (bottom). **(D)** Purified PRC2 or PRC2 in complex with long or short isoform of AEBP2. **(E)** HMT assay containing PRC2 or PRC2 in complex with short or long isoform (15, 30, 60nM) using unmodified di-nucleosomes (200nM) as substrate. The levels of H3K27 methylation are shown by autoradiography (top). Quantification of the relative amount of H3K27 methylation after 60 min of incubation is shown (n = 3/data point) (bottom). **(F)** HMT assay containing PRC2 or PRC2 in complex with AEBP2 short or long isoform (30, 60, 120nM) using H2AK119ub1 mono-nucleosomes (200nM) as substrate. The levels of H3K27 methylation are shown by autoradiography (top). Quantification of the relative amount of H3K27 methylation after 60 min of incubation is shown (n = 3/data point) (bottom).

### The acidic DE-rich motifs exclusively on the N-terminal region of the AEBP2 long isoform are essential for its inhibitory effect on PRC2 activity

We next analyzed the amino acid sequences of AEBP2 isoforms, focusing on the N-terminal region unique to the long isoform, to gain insight into how the long and short isoforms differentially regulate PRC2 activity. The long isoform has an additional N-terminal region with 208 amino acids, including two negatively charged DE-rich motifs (Fig. 2A and S2A).

**Figure 2.**
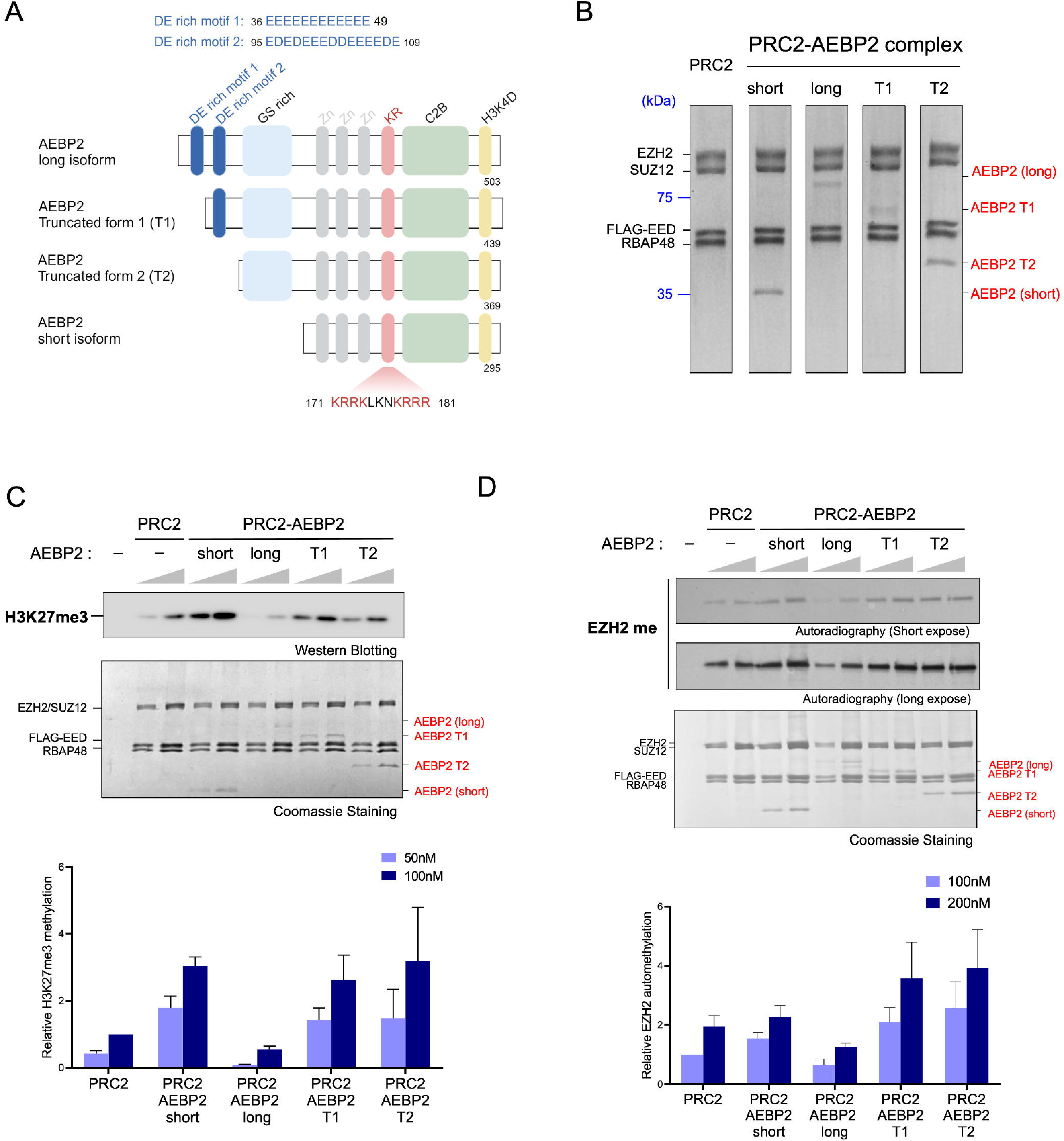
AEBP2 long isoform inhibits PRC2 activity through DE motifs in its N-terminal. **(A)** Schematic representation of full-length AEBP2 short isoform, AEBP2 long isoform, or its truncated AEBP2 fragments (T1 and T2). The KR motif, required for AEBP2 short isoform to enhance nucleosome binding and stimulate PRC2, is indicated. The sequences of DE-rich motifs are shown above. **(B)** Purified PRC2 or PRC2-AEBP2 (left) and purified AEBP2 isoforms and truncated fragments (right). **(C)** HMT assay containing PRC2 or PRC2-AEBP2 complexes (50, 100nM) using di-nucleosomes (150nM) as substrate. The levels of tri-methylation on histone H3 are shown by western blot with H3K27me3 antibody (top). Quantification of the relative amount of H3K27me3 on histone H3K27 after 60 min of incubation is shown (n = 3/data point) (bottom). **(D)** Methyltransferase assay containing PRC2 or PRC2-AEBP2 complexes (50, 100nM) with ^3^H-labeled S-adenosylmethionine (SAM). Autoradiography shows that the long isoform and its DE-rich fragments inhibit EZH2 automethylation. (top). Quantification of the relative amount of EZH2 automethylation after 60 min of incubation is shown (n = 3/data point) (bottom).

Therefore, these motifs are predicted to interact with positively charged regions of PRC2 and influence the PRC2 activity. Our previous studies have shown that the KR motif in AEBP2 is crucial for enhancing nucleosome binding and stimulating PRC2 activity (Lee *et al*, 2018a). Therefore, we hypothesize that these DE-rich motifs may prevent the interaction between the KR motif of AEBP2 and nucleosomal DNA or interact with other positively charged regions within PRC2, ultimately dampening its catalytic activity.

To test whether DE-motifs are involved in PRC2 inhibition, we generated truncation mutants of AEBP2 that lacked either one DE-rich motif (T1) or both DE-rich motifs (T2) (Fig. 2A). We then purified short and long isoforms of AEBP2, along with the T1, and T2 mutants in complex with PRC2 (Fig. 2B). Deletion of one or both DE-rich motifs alleviated the repressive effect of AEBP2 long isoform on PRC2 (Fig. 2C). These data demonstrate that the DE-rich motifs in the N-terminal region of the AEBP2 long isoform contribute to the inhibitory effect on PRC2 activity.

### The DE-rich motif of AEBP2 long isoform minimally influences nucleosome binding while restraining EZH2 automethylation

To test whether DE-motif could interfere with PRC2-AEBP2 and nucleosome binding, we performed an electrophoretic mobility shift assay (EMSA) using either mono-nucleosomes or di-nucleosomes as substrates. Both the long and short forms enhanced the binding of PRC2-AEBP2 to the nucleosome (Fig. S2B and Fig. S2C), likely through the KR-motif which enhance nucleosome binding as previously reported (Lee *et al*, 2018b) (Fig. 2A). Moreover, PRC2-AEBP2 with short isoform showed marginally higher nucleosome binding than that with the long isoform, suggesting that the DE-motif may subtly contact the KR-motif of AEBP2 or engage PRC2 regions that reduced nucleosome binding. We then hypothesized that the N-terminal region of AEBP2 long isoform could directly inhibit the catalytic activity of PRC2. To test this, we conducted methyltransferase assays measuring the EZH2 automethylation level (Lee *et al*, 2019; Wang *et al*, 2019; Popoca & Lee, 2022a). Notably, while the short isoform stimulated EZH2 automethylation, the long isoform inhibited it (Fig. 2D and Fig. S2D), indicating the N-terminal region of the AEBP2 long isoform inhibits the intrinsic catalytic activity of PRC2. AlphaFold modeling revealed that the DE motifs interact with the SET domain of EZH2, potentially occluding the SAM-binding and catalytic sites (Fig. S2E and Fig. S2F), thereby suggesting inhibition of both EZH2 automethylation and H3K27 methylation.

### Distinct chromatin association and modulation of H3K27me3 deposition by AEBP2 isoforms

To determine the chromatin binding of each AEBP2 isoform and H3K27me3 deposition, we performed spike-in normalized ChIP-seq for AEBP2 and spike-in normalized CUT&Tag for H3K27me3 in WT, TKO, and TKO mESCs rescued with either AEBP2 long (AEBP2-L) or AEBP2 short (AEBP2-S) isoform. We found that both isoforms occupy largely overlapping genomic regions, although the long isoform exhibited weaker chromatin association (Fig. 3A). Consistent with western blot results (Fig. 1C), cells rescued with the long isoform displayed lower level of H3K27me3 than those rescued with the short isoform, as reflected in both the global peak intensity and the heatmap distribution (Fig. 3A-3C). Notably, H3K27me3 enrichment at CpG island (CGI) centers was not fully restored even in the short isoform rescue cells (Fig. 3A and Fig. 3B). This incomplete restoration may reflect the requirement for Mtf2 or Jarid2, key cofactors that are required for PRC2 nucleation at CGIs (Oksuz *et al*, 2018). Alternatively, the absence of *Mtf2* and *Jarid2* might lead to increased DNA methylation at CGI cores, thereby limiting PRC2 activity (Yang & Li, 2020; Perino *et al*, 2020). Certain loci, including *Sfi1* and several other regions, displayed even lower H3K27me3 level in the AEBP2 long-isoform rescue than in TKO cells (Fig. S3A), implying that the long isoform suppresses PRC2 activity in cells. Interestingly, the H3K27me3 signal across the *Hoxa* cluster, especially at *Hoxa9*, *Hoxa10* and *Hoxa11* loci, was particularly diminished in the AEBP2 long isoform rescue cells compared with both the WT, TKO and AEBP2 short isoform rescue cells (Fig. 3D). These results suggest that the long AEBP2 isoform selectively attenuates PRC2 activity at lineage-specific developmental loci. Consistently, mRNA expression of *Hoxa9*, *Hoxa10*, *Hoxa11* was increased in AEBP2 long isoform-rescued mESCs relative to WT, TKO and AEBP2 short isoform-rescued cells and this upregulation persisted even after 4 days spontaneous differentiation (Fig. 3E and Fig. S3B).

**Figure 3.**
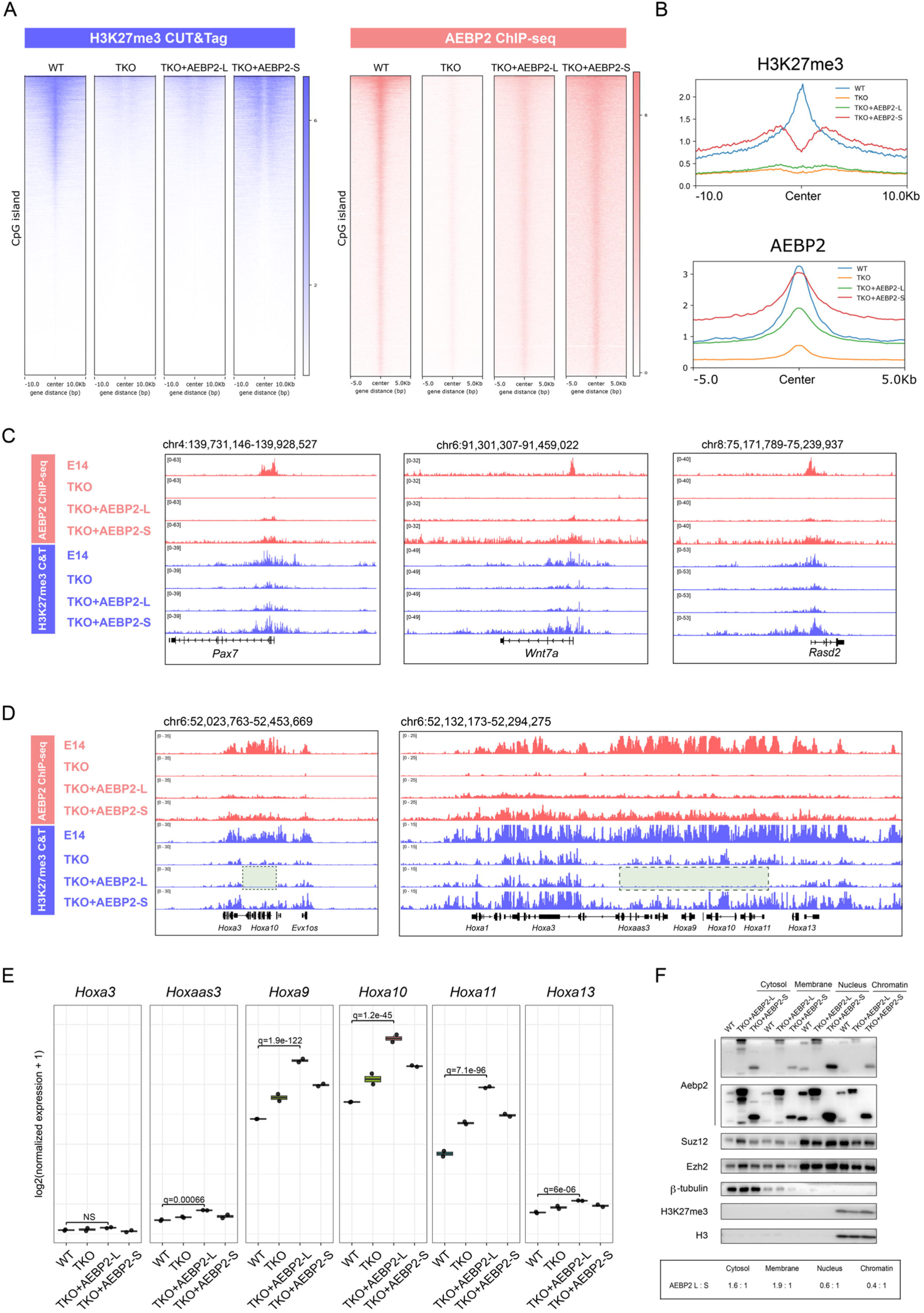
Distinct chromatin association and modulation of H3K27me3 deposition by AEBP2 isoforms. (A) Heatmaps showing spike-in normalized H3K27me3 CUT&Tag (left) and AEBP2 ChIP-seq (right) signals centered by CpG islands (CGIs) in WT, TKO, and TKO mESCs rescued with either AEBP2 long isoform (AEBP2-L) or AEBP2 short isoform (AEBP2-S). **(B)** Average signal intensity profiles of H3K27me3 CUT&Tag and AEBP2 ChIP-seq signals from Panel A. **(C)** Genome browser views showing AEBP2 ChIP-seq (red) and H3K27me3 CUT&Tag (blue) profiles at representative developmental genes (*Pax7*, *Wnt7a*, and *Rasd2*).**(D)** Genome browser views showing AEBP2 ChIP-seq (red) and H3K27me3 CUT&Tag (blue) profiles across the *Hoxa* gene cluster loci. (Right) Average distribution across the *Hoxa* cluster region. (Left) Enlarged view highlighting individual *Hoxa* genes. The green box marks regions exhibiting H3K27me3 depletion specifically in TKO + AEBP2-L cells. (E) Gene expression analysis of *Hoxa* cluster genes during differentiation in AEBP2 isoform–rescued cells. Boxplots showing normalized RNA-seq expression (log₂ fold change; DESeq2 normalized counts) of *Hoxa3*, *Hoxa3as3*, *Hoxa9*, *Hoxa10*, *Hoxa11*, and *Hoxa13* in WT, TKO, and TKO cells rescued with either the AEBP2 long (L) or short (S) isoform. **(F)** Subcellular fractionation showing localization of AEBP2 isoforms in cytoplasmic, nuclear membrane, nucleus, and chromatin-bound fractions.

To further investigate the difference in chromatin recruitment between the AEBP2 long and short isoforms, despite their comparable binding affinity to di-nucleosomes *in vitro* (Fig. S2B and Fig. S2C), we examined their subcellular localization. We fractionated the cell extracts into cytoplasmic, nuclear membrane, nuclear soluble, and chromatin-bound fractions, and analyzed the distribution of each isoform. Notably, the AEBP2 short isoform was more abundant in the nuclear and chromatin fractions, indicating that it is more efficiently retained on chromatin compared to the long isoform (Fig. 3F).

### Lack of AEBP2 short isoform results in impaired early-stage differentiation

To assess the impact of each AEBP2 isoform on early differentiation, we induced spontaneous differentiation of WT, TKO, and TKO cells rescued with either the long (AEBP2-L) or short (AEBP2-S) isoform by withdrawing LIF/2i and monitored the expression of pluripotency markers. Notably, TKO+AEBP2-L cells maintained high levels of Sox2 and Nanog even after 4 days of differentiation (Fig. 4A), indicating that these cells remained in a pluripotent-like state and exhibited impaired early differentiation.

**Figure 4.**
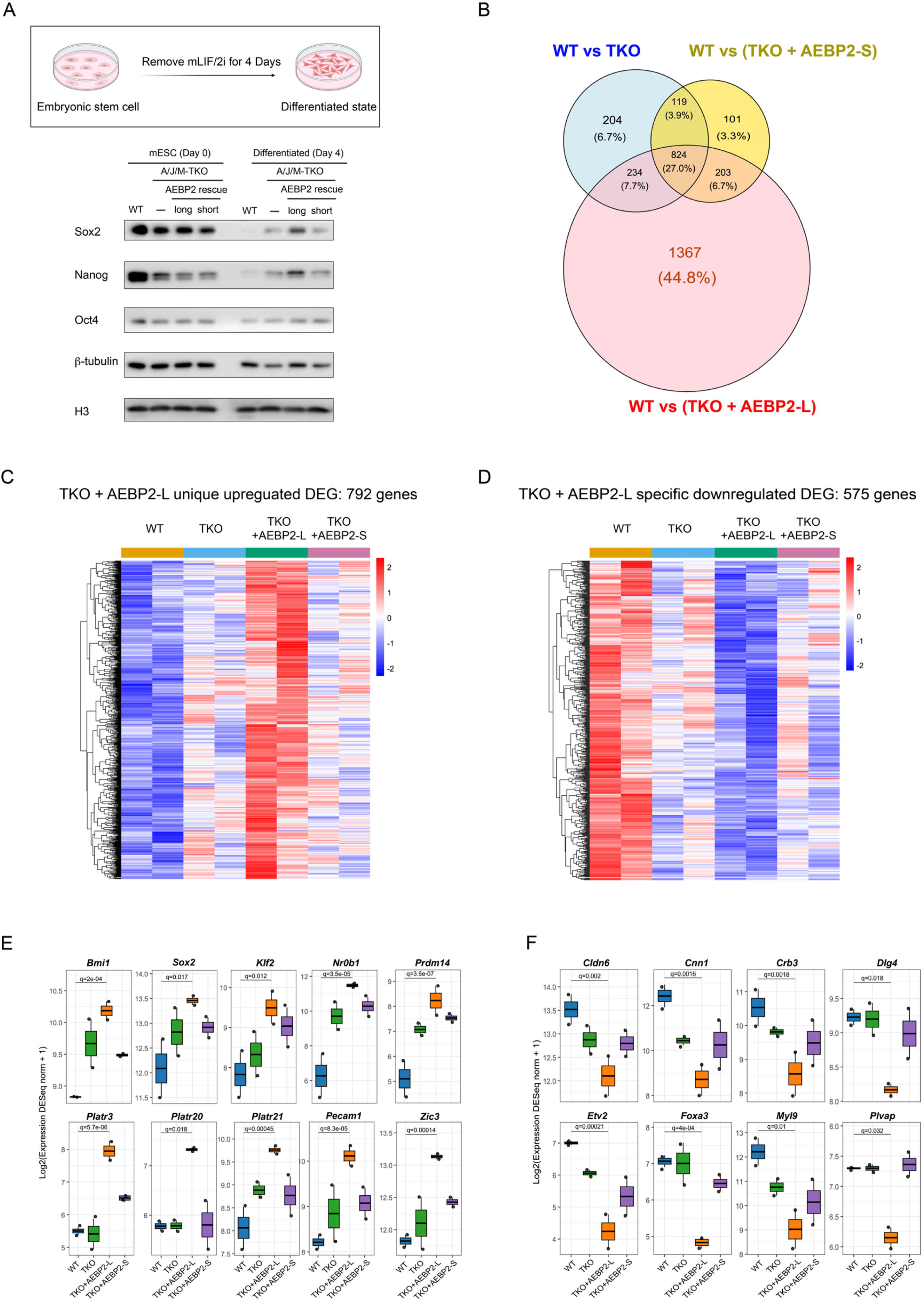
AEBP2 short isoform is crucial for early-stage differentiation from mESCs to embryoid bodies. (A) Schematic representation of spontaneous differentiation of mESCs at day 0 and day 4 following removal of mLIF and 2i (Top). Western blot analysis showing expression levels of pluripotency markers (Sox2, Nanog, Oct4), b-actin and H3 in WT, TKO, and TKO mESCs rescued with either AEBP2 short or long isoform at day 0 and day 4 following spontaneous differentiation (Bottom). b-actin and H3 serve as a loading control. **(B)** Venn diagram showing the overlap of differentially expressed genes (DEGs) between WT versus TKO, WT versus AEBP2 short isoform rescue (TKO+AEBP2-S), and WT versus AEBP2 long isoform rescue (AEBP2-L) cells. The numbers and percentages indicate the number of DEGs unique to each comparison or shared among multiple conditions. **(C-D)** Heatmaps showing representative upregulated (E) and downregulated (F) gene clusters specific to AEBP2 long isoform rescue. **(E-F)** Gene expression analysis of embryonic stem cell-related genes (C) and developmental genes (D) in WT, TKO, and TKO cells rescued with either the AEBP2 long or short isoform. Boxplots showing normalized RNA-seq expression (log₂ fold change; DESeq2 normalized counts) of each gene.

We next performed RNA-seq analysis on differentiated WT, TKO, TKO+AEBP2-L, and TKO+AEBP2-S cells to examine genome-wide transcriptional changes. Among these, the AEBP2 -L rescue showed the largest number of unique differentially expressed genes (1,367), whereas the AEBP2-S rescue exhibited the fewest (101) (Fig.4B). This indicates that the transcriptomic profile of AEBP2-S rescue cells closely resemble that of WT cells after differentiation, in contrast to AEBP2-L rescue cells. Further analysis DEGs specific to AEBP2-L rescue (792 up-regulated and 575 down-regulated genes) revealed increased expression of pluripotency-associated genes, including *Sox2* and *Prdm14*, alongside reduced expression of developmental and lineage-specific genes, such as *Foxa3* and *Dlg4* (Fig 4C-4F).

These findings demonstrate that isoform switching of AEBP2 fine-tunes PRC2 activity to balance repression and differentiation, revealing a regulatory axis that safeguards proper lineage progression.

## Discussion

The activity of Polycomb Repressive Complex 2 (PRC2) is finely tuned by its accessory proteins, whose expression and function vary depending on developmental and cellular context. Most of accessory proteins including PHF1, MTF2, PHF19, JARID2, and the short isoform of AEBP2 stimulate PRC2 activity. In contrast, disease-relevant EZHIP (also known as CATACOMB) antagonizes PRC2 function in a manner similar to H3K27M oncohistone (Piunti *et al*, 2019; Jain *et al*, 2019; Chaouch *et al*, 2021; Stafford *et al*, 2018; Lewis *et al*, 2013).

In this study, we identified the long isoform of AEBP2 as a negative regulator of PRC2 activity. Despite containing a conserved KR motif—shared with the short isoform and known to enhance nucleosome binding—only the long isoform harbors a DE-rich motif, which is essential for its inhibitory function. Deletion of this motif abolishes its repressive activity, indicating that it mediates a specific molecular interaction with PRC2. Structural and biochemical evidence suggests that this DE motif directly engages the SET domain of EZH2, interfering with the SAM-binding or catalytic pocket, thereby repressing PRC2 activity.

This functional duality, KR-mediated chromatin tethering and DE-mediated catalytic inhibition of AEBP2, enables the long isoform to simultaneously recruit PRC2 to chromatin and restrain its catalytic activity. Such architecture likely evolved to provide tighter control over PRC2 activity, particularly in contexts where widespread H3K27 methylation must be restricted.

Consistent with this model, the expression of AEBP2 isoforms is dynamically regulated across developmental stages. The short isoform of AEBP2 is predominantly expressed during early development, coinciding with high levels of PRC2 core components such as EZH2, EED, and SUZ12. During this stage, as cells undergo rapid proliferation while preserving pluripotency, robust PRC2 catalytic activity is essential to sustain H3K27 methylation following DNA replication. The abundant expression of the AEBP2 short isoform likely plays a key role in enhancing PRC2 activity to sustain this biological requirement, highlighting a functional correlation between isoform expression and PRC2 catalytic activity. By contrast, the requirement for PRC2 activity decreases in terminally differentiated, non-dividing cells. This reduction is accompanied by the downregulation of PRC2 core components and a parallel decline in the expression of its accessory proteins (Lee *et al*, 2018b; Benoit *et al*, 2012; Vizán *et al*, 2020; Zhang *et al*, 2011a). Notably, differentiated cells exhibit a marked loss of the AEBP2 short isoform, with sustained expression of the long isoform as the predominant variant in differentiated cells. This isoform switch, from an activator to an inhibitor of PRC2, reflects the reduced catalytic needs of the complex, suggesting that developmental stage-specific expression of AEBP2 isoforms provides a mechanism to dynamically modulate PRC2 activity according to cell state.

Mechanistically, AEBP2 may coordinate with other PRC2.2 components, particularly JARID2, to regulate H3K27 methylation dynamics. While MTF2 (a PRC2.1 subunit) anchors PRC2 at CpG islands and maintains repression, JARID2 promotes *de novo* recruitment during early differentiation. AEBP2 and MTF2 compete for SUZ12 binding, forming mutually exclusive PRC2.2 and PRC2.1 complexes. As PRC2 transitions from initiation sites into broader chromatin domains, a switch from PRC2.1 to PRC2.2 may occur, with AEBP2 facilitating spreading or repression depending on the isoform expressed. Notably, JARID2 can allosterically activate PRC2, and the presence of the DE motif in AEBP2 long isoform may serve as a counterbalance, acting as a molecular gatekeeper to prevent hyperactivation.

Although JARID2 is a critical PRC2 accessory factor with established roles in recruitment, activity, and spreading, it was not included in the current analysis due to its structural complexity, which could confound the mechanistic dissection of AEBP2 function. Future studies are needed to elucidate the combinatorial effects of AEBP2 and JARID2 and how their interactions govern PRC2 behavior under physiological conditions.

Finally, our findings emphasize the importance of isoform-specific regulation in chromatin control. DE-rich motifs—clusters of acidic residues such as aspartic acid (D) and glutamic acid (E)—are well known to mediate electrostatic interactions, as seen in nucleosome acidic patches (Leung *et al*, 2014; Baier *et al*, 2024; Liu *et al*, 2023) and PRC2 subunit interfaces (Grau *et al*, 2021). These motifs modulate protein-protein interactions, chromatin compaction (Kalashnikova *et al*, 2013), and enzyme regulation. The DE motif in AEBP2 exemplifies how such domains can evolve to impose precise functional control within chromatin-modifying complexes.

In summary, the antagonistic roles of AEBP2 isoforms reveal a finely balanced regulatory system through which isoforms encoded by the same gene can exert both activating and repressive effects on PRC2. This isoform-based regulation enables context-dependent tuning of PRC2 activity during development and differentiation, adding to the growing framework that alternative splicing and modular domain architecture are key strategies for achieving epigenetic precision.

## Data availability

This paper does not report original code.

Any additional information required to reanalyze the data reported in this paper is available from the lead contact upon request.

## Author contributions

Y.L. and C-H.L. conceptualized and designed the study. Y.L. and H.K. conducted the experiments and bioinformatics analyses. Y.L. and C-H.L wrote the manuscript.

## Acknowledgements

We thank Drs. Jia-Ray Yu and Karim-Jean Armache for critical reading of the manuscript as well as Lee laboratory members for critical comments and discussions. This study was supported by the Korean government (MSIT) (grant numbers RS-2023-00223069, RS-2025-02217909, RS-2025-24523684, RS-2022-NR070833 and NRF2021R1C1C1013220) and the BK21 Four Biomedical Science Program. The SNUH Kun-hee Lee Child Cancer and Rare Disease Project Foundation, Republic of Korea (grant number 22B-001-0100), the Research Resettlement Fund for the new faculty of Seoul National University, the Creative-Pioneering Researchers Program through Seoul National University, grants from Seoul National University College of Medicine, the AI-Bio Research Grant through Seoul National University also supported this study. Furthermore, it was supported by Doosan Yonkang Foundation (Grant No. SNUH-30-2024-0440) and by NAVER (Grant No. SNUH-3720242170). The figures were created with BioRender.com.

## Materials and Methods

### Cell culture

Mouse embryonic stem cells (E14 and OG2) and derivates were grown on 0.5% gelatin-coated dishes in DMEM containing 15% (v/v) fetal bovine serum, Penicillin streptomycin (10000U/mL), 2mM L-Glutamine, 1X MEM Non-Essential Amino Acids, 10mM 2-Mercaptoethanol and mouse Leukemia Inhibitory Factor (LIF), 1 mM MEK1/2 inhibitor (PD0325901) and 3 mM GSK3 inhibitor (CHIR99021) at 37°C with 5% CO_2_.

### Recombinant bacterial protein

Human AEBP2 long (isoform 1 in human, a.a. 1∼503), short (a.a. 219∼503) and truncated forms (T1, a.a 75∼503, and T2, a.a 145∼503) were cloned into pET21a. Strep-tagged AEBP2 long, short and truncated forms were expressed in *E. coli* strain Rosetta with LB Media for 16 h at 18°C by induction with 0.5 mM Isopropyl β-D-1-thiogalactopyranoside (IPTG). E. coli cells were resuspended in 20 mM Tris-HCl, pH 7.5, 350 mM NaCl, 10% glycerol and protease inhibitor (1 mM phenylmethlysulfonyl fluoride (PMSF), 1 mM benzamidine, 1μg/ml Leupeptin and 1μg/ml pepstatin). Cell lysate purified using Streptactin Sepharose beads (Cytiva) for 3 h at 4°C and eluted by desbiotin and the proteins were again purified with Q Sepharose beads (Cytiva). Purified proteins were dialyzed with 20 mM Tris-HCl, pH 7.5, 350 mM NaCl and 10% glycerol.

### Protein purification using baculovirus expression system

Baculovirus expression plasmids for human EZH2, SUZ12, Flag-tagged EED and RBAP48 were cloned into pFastBac1. The recombinant pFastBac1 vectors were transformed to *E. coli* DH10Bac cells to create recombinant bacmid. Recombinant human PRC2 core subunits (EZH2, EED, SUZ12, and RBAP48) were co-expressed in Sf9 insect cells grown in SFM Media (Expression System). After 64 hrs of infection, cells were resuspended in 20 mM Hepes-HCl, pH 7.5, 350 mM NaCl, 10% glycerol, and protease inhibitor (see above) and lysed with a cell disruptor (machine name). Then PRC2 complex was purified using FLAG-M2 agarose beads (Sigma) and Q Sepharose (Cytiva). Purified proteins were dialyzed with 20 mM Tris-HCl, pH 7.5, 180 mM NaCl and 10% glycerol. For generating PRC2-AEBP2 complexes, AEBP2 long, short, and truncated forms purified from the *E. coli* system were incubated with PRC2 Sf9 cell lysate before purification. The purification step was same as PRC2 core complex purification mentioned above.

### Nucleosome reconstitution

Recombinant histones were generated as previously described(Luger *et al*, 1997). Briefly, each histone was expressed in Rosetta (DE3) cells (Novagen), extracted from inclusion bodies, and purified by sequential anion chromatography and size exclusion chromatography. For refolding recombinant octamers, equimolar amounts of histones were mixed and dialyzed into refolding buffer (10 mM Tris-HCl, pH 7.5, 2 M NaCl, 1 mM EDTA, and 5 mM β-mercaptoethanol). Octamers were further purified by size exclusion chromatography on a Superdex 200 column (GE healthcare) in refolding buffer. Recombinant oligonucleosomes were reconstituted by gradient salt dialysis of octamers and indicated DNA fragments or Cy5-labeled 601-nucleosome positioning sequences.

### HMT assay

Standard HMT assays were performed in a total volume of 15mL containing HMT buffer (50 mM Tris-HCl, pH 8.5, 5 mM MgCl_2_, and 4 mM DTT) with either 50mM of cold S-Adenosylmethionine (SAM, Merck) or 1uM Adenosyl-L-Methionine, S-[methyl-3H]-(Revvity), 100nM of nucleosomes, and recombinant human PRC2 complexes under the following conditions. The reaction mixture was incubated for 60 min at 30°C and stopped by adding 4 μL of STOP buffer (0.2 M Tris-HCl, pH 6.8, 20% glycerol, 10% SDS, 10 mM β-mercaptoethanol, and 0.05% Bromophenol blue). The yield of each HMT reaction was measured as follows:

After the HMT reactions, samples were incubated at 95°C for 5 minutes and then separated using SDS-PAGE. Proteins were transferred to 0.45 µm PVDF membranes (Millipore) via wet transfer. For reactions using cold SAM, the H3K27me3 levels were detected by Western blot using an H3K27me3-specific antibody. For reactions using hot SAM, the total levels of H3K27 methylation and EZH2 methylation were assessed by autoradiography(Popoca & Lee, 2022b).

### Gel mobility shift assay

Cy5-labeled core nucleosomes were incubated with PRC2 or PRC2-AEBP2 complexes in 10 mM HEPES, pH 7.9, 50 mM KCl, 1 mM DTT and 5% glycerol in a total volume of 15 μL for 30 min at room temperature. Complex formation was analyzed by 4% native polyacrylamide gel electrophoreses (0.3 x TBE, 5% glycerol). Cy5 signal was detected using the ChemiDoc™ MP Imaging System (Bio-Rad Laboratories, USA).

### Preparation of whole cell extract and Western-Blotting

Cells were harvested and lysed with RIPA buffer (10mM Tris-HCL, pH7.5, 400mM NaCl, 1mM EDTA, 1% Trition X-100, 0.1% sodium deoxycholate, and 0.1% SDS) containing protease inhibitors (0.5mM PMSF, 1mM Benzamidine-HCl, 1μg/mL Pepstatin, 1μg/mL Leupeptin) and phosphatase inhibitors (10 mM NaF and 1 mM Na3VO_4_). The cell suspension was sonicated (40% amplitude, 12 strokes) and centrifuged at 20,000 x g at 4°C for 10 min. The supernatant (whole cell extract) was mixed with denaturing sample buffer, boiled for 5 min and run onto 4-16% SDS-PAGE gels. The gels were transferred on PVDF membrane at 100V for 90min. PDVF is blocked in TBS-T-Milk for 1h at RT, then O/N at 4°C with primary antibody and 1h at RT with secondary antibody.

### Lentiviral production and delivery

AEBP2 short isoform, long isoform and truncated fragments of long isoform (T1 and T2, respectively) were subcloned into the pLV-EF1-alpha-IRES-Pruo vector (Clontech). Subcloned lentiviral vectors were co-transfected with pcREV, BH-10, and pVSV-G packaging vectors (Viral vector 10 mg, VSVG 3 mg, BH10 5 mg, and pcREV 2.5 mg) into Lenti-X 293T cells to produce viral particles. Virus-containing medium was collected 48 h after transfection and polybrene was added to the viral medium at a concentration of 8 μg/ml. The target cells were infected and sorted for mCherry positive by FACS cytometry (BD FACSAria™ III).

### Generation of CRISPR/Cas9-mediated genome editing cell line(s)

Aebp2 KO cell line was obtained from previous study(Lee *et al*, 2018b) The *Aebp2, Mtf2, and Jarid2* triple-knockout (TKO) cell line was generated by introducing Mtf2 and Jarid2 knockouts into the Aebp2 KO background. The gRNA target sequences were designed using the http://www.rgenome.net/cas-offinder/ design tool. The sgRNA was cloned into the SpCas9-2A Puro (Addgene: PX459) via BbsI digestion and insertion. mESC cells were seeded into 6-well plates at 80,000 cells per well and transfected with 1.5μg of the appropriate guide RNAs, template DNA for guide RNAs and Cas9 endonuclease using Lipofectamine 2000 (Thermofisher). The transfection was performed using a 2:1 ratio of Lipofectamine: DNA. After transfection, cells were selected with 2 μg/ml puromycin for 48 h and single ESC clones were picked after 7-10 days, trypsinized in 0.25% Trypsin-EDTA for 5 min, and plated into individual wells of a 96-well plate for genotyping. Each clone was characterized to confirm gene knockout by Western-Blot and gDNA sequencing.

### CUT&Tag

Cells were collected and washed twice with PBS, then fixed with 0.1% formaldehyde for 1 minute. The fixation was quenched by adding 0.125 M glycine, followed by another PBS wash. The cells were resuspended in Nuclear Extraction (NE) buffer (20 mM HEPES, pH 7.9, 10 mM KCl, 0.1% Triton X-100, and 20% glycerol) and incubated on ice for 10 minutes. Following nuclear extraction, the nuclei were washed with cold NE buffer and resuspended in cold NE buffer at a concentration of 0.05 million nuclei/100 µL. The nuclei were then immobilized on Concanavalin A Conjugated Paramagnetic Beads (Epicypher, 21-1401). After immobilization, the nuclei were permeabilized with 0.01% digitonin and incubated overnight at 4°C with the primary antibody at the manufacturer’s recommended concentration. The CUT&Tag-IT Spike-In Control (Active Motif, 53168) was included in parallel following the manufacturer’s protocol to allow normalization of signal intensity across samples. Afterward, samples were incubated with 0.5 µg of secondary antibody for 1 hour. Following the secondary antibody incubation, pA-Tn5 (Epicypher, 15-1017) was added, and the samples were incubated for 1 hour at room temperature. Targeted tagmentation was mediated by adding magnesium chloride (10mM MgCl2) and the reactions were incubated at 37°C for 1 hour. After tagmentation, the reaction was stopped, and the DNA fragments were amplified using PCR. The libraries were quantified, quality-checked, and sequenced on an Illumina NovaSeq X Plus platform using 150 bp paired-end reads.

### ChIP-seq

For double crosslinking, 3 × 10^6^ cells were harvested and fixed sequentially with 2mM disuccinimidyl glutarate (DSG; Thermo Fisher, 20593) for 35 minutes followed by 1% formaldehyde for 10 minutes at room temperature. Crosslinking was quenched with 0.125M glycine, and nuclei were isolated in NE buffer and extracted nuclei were then resuspended 1% SDS lysis buffer (50mM Tris-HCl pH 8, 10mM EDTA pH 8, 1% SDS) and sonicated by Covaris S220 sonicator (Power 175 W, Duty 10%, 200 cycles per burst) for 430 seconds to generate DNA fragments of approximately 200bp. Sheared chromatin was diluted in ChIP dilution buffer (16.7 mM Tris-HCl, pH 8.0, 0.01% SDS, 1.1% Triton X-100, 16.7 mM NaCl, 1.2 mM EDTA) and clarified by centrifugation at 13,000 rpm for 15 minutes at 4 °C.

Chromatin was incubated overnight at 4 °C with Dynabeads protein A pre-bound to 2.4mg target antibody. 1 µg of Drosophila chromatin (Active Motif, #53083) and 0.1 µg of anti-Drosophila H2A.X antibody (Active Motif, #61686) were added per reaction to perform spike-in normalization. The next day, Dynabeads were washed three times with buffer A (140mM NaCl, 1mM EDTA pH 8, 0.5mM EGTA pH 8, 1% Triton X-100, 0.1% SDS, 0.1% sodium deoxycholate, 10mM Tris-HCl pH 8), twice with buffer B (300mM NaCl, 1mM EDTA pH 8, 0.5mM EGTA pH 8, 1% Triton X-100, 0.2% SDS, 0.1% sodium deoxycholate, 10mM Tris-HCl pH 8) and once with buffer C (250 mM LiCl, 10mM Tris-HCl pH 8, 1mM EDTA, 0.5% NP-40, 0.5% sodium deoxycholate). RNA was removed by treatment with RNAse A at 37 °C for 30 minutes and crosslinks were reversed overnight at 65 °C in the presence of Proteinase K and SDS treatment.

Eluted DNAs were size selected using SPRIselect beads (Beckman B23319) and Libraries were prepared with NEB DNA library prep kit (NEB, E7645L) following the manufacturer’s protocol. The libraries were quantified, quality-checked, and sequenced on an Illumina NovaSeq X Plus platform using 150 bp paired-end reads.

### ChIP-seq and CUT&Tag analysis

Adaptor sequences were trimmed using Cutadapt, and the filtered reads were aligned to the mouse reference genome (mm10) with Bowtie2 using default parameters. Duplicate reads were identified and removed with Samtools markdup. Reads that failed to align to the mouse genome were subsequently mapped to the Drosophila genome (dm6), and the number of Drosophila-mapped reads was used to calculate the spike-in normalization factor. Normalized bigWig files, heatmaps, and average profiles were generated using deepTools.

### RNA-seq

A total of 5 × 10⁶ cells were harvested and resuspended in 1 mL of TRIzol reagent (Thermo Fisher, #15596026), followed by incubation for 5 min at room temperature. Chloroform (0.2 mL) was added, and the samples were vortexed vigorously and centrifuged at 13,000 rpm for 15 min at 4 °C. The upper aqueous phase containing RNA was transferred to a new tube, and RNA was precipitated with isopropanol and washed with ice-cold 70% ethanol. The RNA pellet was air-dried, dissolved in DEPC-treated water (Invitrogen, #AM9915G), and its concentration and integrity were assessed. Sequencing libraries were prepared using the TruSeq Stranded Total RNA Ribo-Zero Gold Kit (Human/Mouse/Rat) (Illumina, #RS-122-2301). The libraries were then sequenced on an Illumina NovaSeq X Plus platform to generate 150 bp paired-end reads.

### RNA-seq analysis

Adaptor sequences were trimmed using Trimmomatic, and the filtered reads were aligned to the mouse reference genome (mm10) using HISAT2. Transcript assembly and quantification were performed using StringTie, and gene-level read counts were obtained with featureCounts. Genes with zero total counts across all samples were removed prior to downstream analyses. Normalization factors were estimated using the median-of-ratios method implemented in DESeq2, and normalized expression values were calculated as log₂(DESeq2-normalized counts + 1). Differential expression analysis was carried out using DESeq2 (Wald test), and genes with an adjusted p-value < 0.05 and |log₂ fold change| > 1 were considered significantly differentially expressed. Gene Ontology (GO) and pathway enrichment analyses were performed using the clusterProfiler R package. Visualization of normalized expression levels and heatmaps was conducted in R (v4.2.3) using ggplot2 and pheatmap.

### mESC differentiation

Approximately 1 × 10⁵ mESCs were plated on 60-mm dishes, and differentiation was induced by withdrawal of LIF, PD0325901, and CHIR99021 from complete medium. Medium was replaced daily, and cells were harvested after 4 days of LIF/2i removal.

### Subcellular protein fractionation

Subcellular protein fractionation was performed using the Subcellular Protein Fractionation Kit for Cultured Cells (Thermo Fisher, #78840) according to the manufacturer’s protocol. The resulting cytoplasmic, membrane, nuclear, and chromatin fractions were analyzed by western blotting using standard procedures.

