## Supplemental figures for "Distinct regulation of AEBP2 isoforms on PRC2 activity"

Supplementary Figure 1

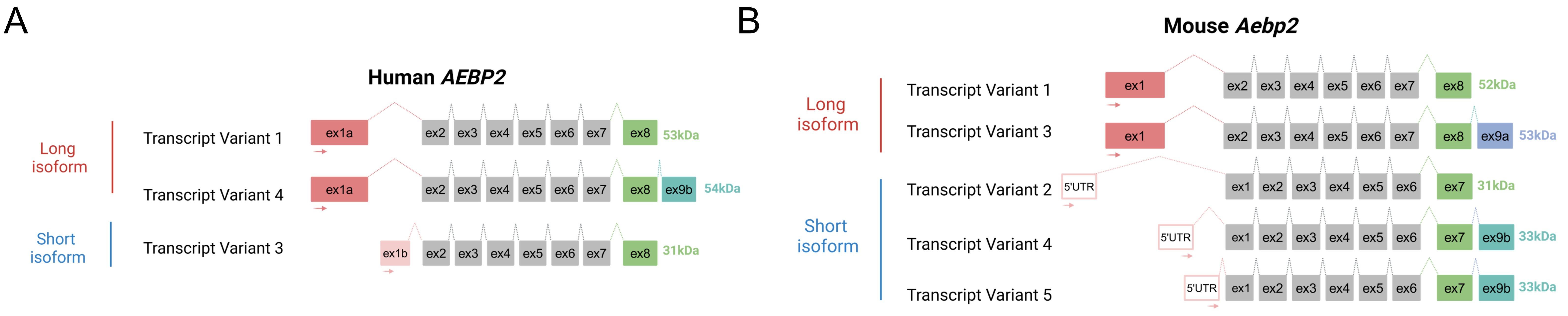


C

```
                       ex1a
Human AEBP2 Long    MAAAITDMADLEELSRLSPLPPGSPGSAARGRAEPPEEEEEEEEEEEEEAEAEAVAALLLN  58
Mouse Aebp2 Long    MAAALADMADLEELSRLSPLSPGSPGPAARGRAEPPEEEEEEED--DEEAEAEAVAALLLN  60
Human AEBP2 Short    -------------------------------------------------------------  0
Mouse Aebp2 Short    -------------------------------------------------------------  0

Human AEBP2 Long    GGSGGGGGGGGGGVGGGEAETMSEPSPESASQAGEDEDEEEDDEEEEDESSSSGGGEEES  120
Mouse Aebp2 Long    GGAGG-------GAGGGEAETMSEPSPESASQAGGDEDEDEED--DEDEGSSSGGAEEES  109
Human AEBP2 Short    ------------------------------------------------------------  0
Mouse Aebp2 Short    ------------------------------------------------------------  0

Human AEBP2 Long    SAESLVGSSSGGCSGDETRSLSPGAASSSSGDGDGKEGLEEPKGPRGGPGGPGSSGGGSS  176
Mouse Aebp2 Long    SAESLVGSSSGGCSGDETRSLSPGAASSSSGDGDGKEGLEEPKGPRGGPGGPGSSGGGSS  169
Human AEBP2 Short    ------------------------------------------------------------  0
Mouse Aebp2 Short    ------------------------------------------------------------  0

Human AEBP2 Long    SSSVVSSGGDEGYGTGGGGSSATSGGRRGSLEMSSDGEPLSRMDSEDSISSTIMDVDSTI  236
Mouse Aebp2 Long    SSSVVSSGGDEGYGTGGGGSSATSGGRRGSLEMSSDGEPLSRMDSEDSISSTLMDIDSTI  229
Human AEBP2 Short    ---------------------------------------MYTRRYSSISSTIMDVDSTI  20
Mouse Aebp2 Short    -----------------------------------------------------MDIDSTI  7
                                                          ex1b
                                       .
                                       .
                                       .
                                         ex9b
Human AEBP2 Long    LSKLPKDTALLLDPNIYRTMPQKRLKRTLIRKVFNLYLSKQ  517
Mouse Aebp2 Long    LSKLPKDTALLLDPNIYRTMPQKRLKRFDILNFPR------  504  ex9a
Human AEBP2 Short    LSKLPKDTALLLDPNIYRTMPQKRLKRTLIRKVFNLYLSKQ  301
Mouse Aebp2 Short    LSKLPKDTALLLDPNIYRTMPQKRLKRTLIRKMFNLYLSKQ  288
```

### Supplementary Figure 2

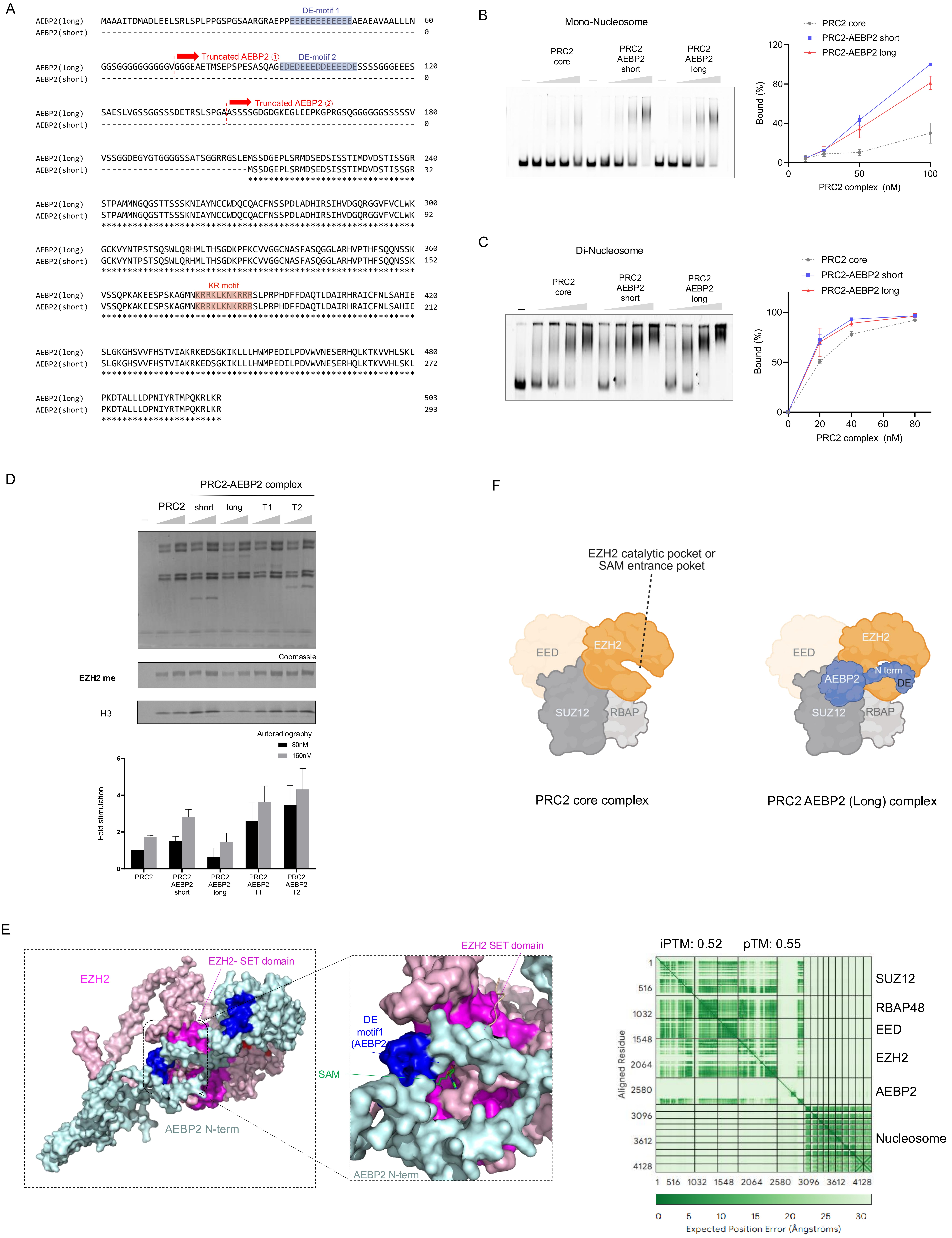

Supplementary Figure 3

A

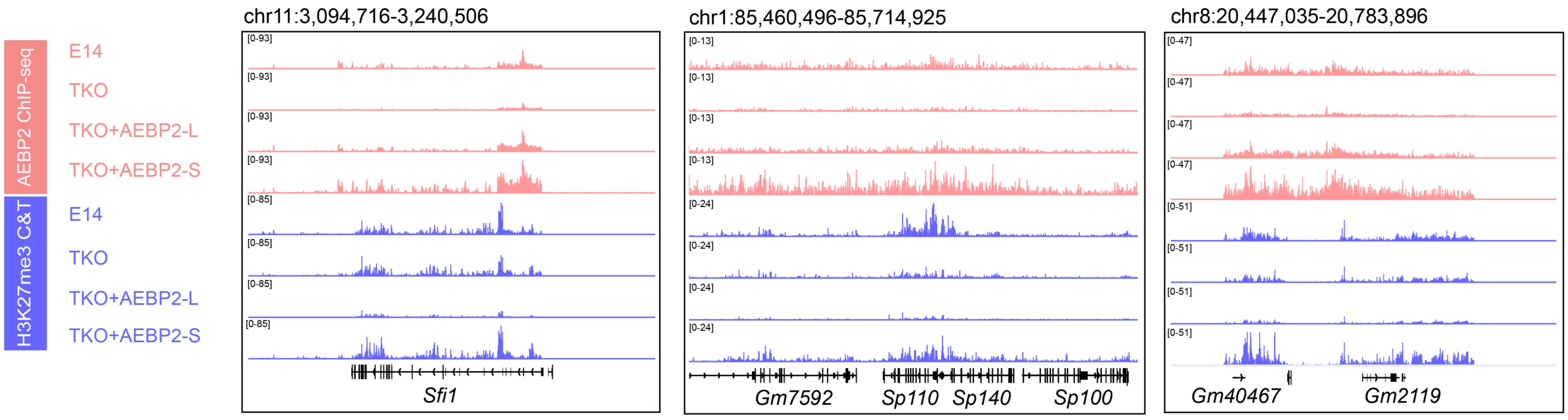


B

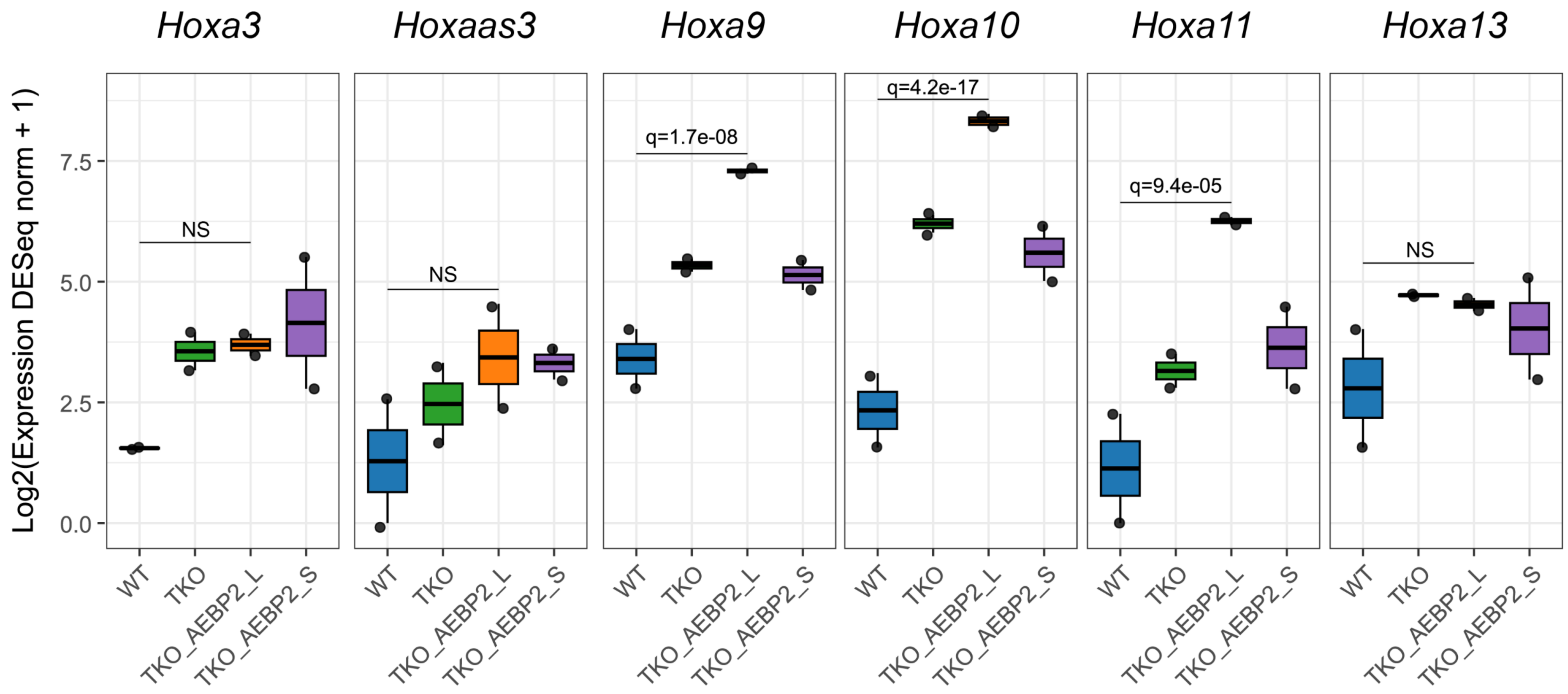
